# Restoration of circulating 2-arachidonoylglycerol levels attenuates cocaine self-administration through endocannabinoid–dopamine interactions in common marmosets

**DOI:** 10.64898/2026.09.01.748459

**Authors:** Chul Kyu Lee, Heejong Eom, Taeyun Yoo, Sun Mi Gu, Jae Jun Lee, Hyeokjun Kwon, Sang-Beom Na, Eunbin Kim, Tae Hwan Kim, Chun-Woong Park, Bang Yeon Hwang, Seong Shoon Yoon, Dohyun Lee, Jaesuk Yun

## Abstract

To date, no pharmacotherapy has been approved for cocaine use disorder. Endocannabinoid signaling is closely related to dopamine-dependent reinforcement and may regulate cocaine-related behaviors. In this study, we used common marmosets (*Callithrix jacchus*) to investigate whether 2-arachidonoylglycerol (2-AG), an endogenous cannabinoid lipid, reduces cocaine reinforcement in a non-restraint oral self-administration model. Marmosets performed oral cocaine self-administration under a fixed-ratio 1 schedule, and cocaine intake was confirmed using plasma benzoylecgonine detection. Dopamine transporter (DAT)-related positron emission tomography (PET) signal, DAT and G protein-coupled receptor 55 (GPR55)- associated fluorescent signals, GPR55/DAT immunofluorescence, and synaptosomal dopamine responses were assessed using 18F-N-(3-fluoropropyl)-2β-carboxymethoxy-3β-(4-iodophenyl) nortropane (^18^F-FP-CIT) PET imaging, ex vivo fluorescent ligand imaging, confocal microscopy, and dopamine quantification. Marmosets acquired oral cocaine self-administration with preferential active lever responding. Repeated cocaine self-administration reduced striatal DAT-related ^18^F-FP-CIT PET signal in vivo. In striatal slices, cocaine decreased fluorescent false neurotransmitter (FFN102)-associated DAT signal and increased T1117-associated GPR55 signal. Systemic 2-AG pretreatment reduced cocaine self-administration while restoring diminished circulating 2-AG levels. GPR55 and DAT immunofluorescent signals were colocalized in marmoset brain sections, and 2-AG enhanced cocaine-induced dopamine elevation in synaptosomal preparations. Taken together, these findings suggest that 2-AG attenuates cocaine-taking behavior and may be associated with GPR55/DAT-related dopaminergic responses in common marmosets.

## 1. Introduction

Cocaine use disorder remains a major public health issue because relapse is common and no approved pharmacotherapy is available. Cocaine blocks dopamine reuptake, increases extracellular dopamine levels, and produces behavioral changes such as hyperactivity, stereotyped movements, and compulsive drug-taking behavior (Hummel and Unterwald, 2002). Endocannabinoid signaling is linked to brain structure and function (Manza *et al*., 2020), and cocaine-induced behaviors are partly regulated by endocannabinoid-related mechanisms, including cannabinoid receptor type 1 (CB1), anandamide, and fatty acid amide hydrolase in reward-related brain regions (De Sa Nogueira *et al*., 2022, Zapata and Lupica, 2021, Scherma *et al*., 2019, Chauvet *et al*., 2014, Rivera *et al*., 2013b). These findings suggest that endocannabinoid signaling may be involved in cocaine reinforcement and cocaine-related behavior.

Non-human primates, especially squirrel monkeys and rhesus monkeys, have been widely used to study cocaine self-administration and drug reinforcement (Justinova *et al*., 2015, Porrino *et al*., 2016). The common marmoset (*Callithrix jacchus*) is a useful small primate model because it has human-like neuroanatomical features, complex behavior, and efficient reproduction and can be handled relatively easily (Nuara *et al*., 2022, Bakken *et al*., 2021, Davis *et al*., 2020, Marx, 2016, Kishi *et al*., 2014, Mitchell and Leopold, 2015, Banks *et al*., 2017). Previous studies have examined cocaine-induced behavioral changes in marmosets using conditioned place preference, figure-eight maze, locomotor activity, and hypervigilance-related behavioral tests (Frankowska *et al*., 2021, De Souza Silva *et al*., 2006, Cagni *et al*., 2012). However, marmosets have been used less often to study endocannabinoid regulation of voluntary cocaine-taking behavior.

The circulating endocannabinoid system is altered in substance use conditions. Plasma concentrations of endocannabinoids, including 2-arachidonoylglycerol (2-AG) and 2-linoleoylglycerol (2-LG), are reduced in abstinent individuals with cocaine use disorder (Pavón *et al*., 2013, Pedraz *et al*., 2015). In animal models, drug self-administration changes brain endocannabinoid concentrations (Caillé *et al*., 2007). Although CB1 blockade has been considered for cocaine use disorder (Martín-García *et al*., 2016), the CB1 inverse agonist rimonabant was withdrawn because of related psychiatric adverse effects (Nguyen *et al*., 2019). Therefore, endogenous cannabinoid lipids and non-CB1 targets may offer alternative ways to regulate cocaine-related behavior. Among these, 2-AG is a full agonist of CB1 and CB2 receptors (Lu *et al*., 2019, Bie *et al*., 2018). Furthermore, endocannabinoids act through other targets, including G protein-coupled receptor 55 (GPR55), serotonin-related pathways, dopamine systems, and transient receptor potential vanilloid 1 (TRPV1) (Oakes *et al*., 2019, Haj-Dahmane and Shen, 2011, Khansari *et al*., 2020). Increasing endocannabinoid signaling reduces relapse-like cocaine-seeking behavior through TRPV1-mediated glutamate homeostasis (Zhang *et al*., 2021), and GPR55 is related to drug reward-related behaviors (Liu *et al*., 2021, Sánchez-Zavaleta *et al*., 2023, Xi *et al*., 2024). These findings suggest that 2-AG may regulate cocaine reinforcement through interactions with dopamine-related synaptic mechanisms.

In the present study, we aimed to determine whether 2-AG modulates cocaine reinforcement and GPR55/DAT-associated dopaminergic signaling in common marmosets.

## 2. Materials and Methods

### 2.1 Animals

Common marmosets (*Callithrix jacchus*) were supplied by the Osong Medical Innovation Foundation (Chungbuk, Republic of Korea). Animals were housed under the controlled conditions of 27 ± 2 °C, 40 ± 10% relative humidity, a 12 h/12 h light/dark cycle (lights on 07:00–19:00; illumination ≥500 Lux), and 10–15 air changes per hour. Marmosets were provided a commercial marmoset diet (50 g/day; No. 0630, Altromin Spezialfutter GmbH & Co. KG, Lage, Germany) and sterilized tap water ad libitum. Environmental enrichment, including wooden perches, beds, and nest boxes, was provided to minimize stress.

For cocaine self-administration and endocannabinoid pharmacology experiments, eleven common marmosets (nine males and two females; 247–400 g) were used. Four marmosets were used for food training, oral cocaine self-administration, plasma benzoylecgonine analysis, and 18F-N-(3-fluoropropyl)-2β-carboxymethoxy-3β-(4-iodophenyl) nortropane ^(18^F-FP-CIT PET) imaging. Four drug-naïve marmosets were used for ex-vivo brain slice imaging and tissue-based assays. Three marmosets were used for cocaine self-administration experiments with vehicle or 2-AG pretreatment and plasma 2-AG measurement. These experiments were approved by the Institutional Animal Care and Use Committee of the Osong Medical Innovation Foundation (KBIO-IACUC-2020-148).

### 2.2 Drugs and reagents

For oral cocaine self-administration, cocaine hydrochloride (MacFarlan Smith Ltd., Edinburgh, UK) was dissolved and delivered orally at 0.4 mg/mL, with 0.1 mL per active lever response. The dose was selected based on previous studies on non-human primate self-administration (Meisch, 2001, Meisch and Stewart, 1995, Carroll *et al*., 2016).

2-AG (Cayman Chemical, Ann Arbor, MI, USA) was dissolved in a vehicle containing 2.5% dimethyl sulfoxide (DMSO) and 2.5% Tween 80 in saline and administered intraperitoneally at 0.01 or 0.1 mg/kg 1 h before cocaine self-administration. Because endocannabinoid lipids are chemically labile, 2-AG–containing solutions were freshly prepared, protected from light, kept on ice when necessary, and used immediately after preparation.

### 2.3 Non-restraint oral cocaine self-administration

The operant chamber was designed and manufactured in-house with modifications from a Med Associates system (Med Associates Inc., Georgia, VT, USA). The chamber measured 44 cm (W) × 74.5 cm (L) × 64 cm (H) externally and 40 cm (W) × 35 cm (L) × 42 cm (H) internally. Self-administration was performed under a fixed-ratio 1 (FR1) schedule in non-restrained, freely moving marmosets.

During food training, marmosets received 50% dextrose syrup after active lever pressing. Animals that produced more than 40 lever responses during food training were advanced to cocaine self-administration. In cocaine self-administration sessions, pressing the active lever resulted in oral cocaine delivery (0.4 mg/mL, 0.1 mL/response), whereas pressing the inactive lever resulted in saline delivery. Each active lever response was followed by a 5-s red cue light and a 5-s timeout. Dextrose syrup or cocaine solution was delivered through a spout for 1 s.

Food training comprised 3-h sessions, 4–5 days per week, for 4–12 weeks until stable responding was established. Stable responding was defined as a pattern in which the number of obtained rewards across three consecutive sessions did not vary by > 25% of the mean number of rewards (Kohut and Bergman, 2016, Mello *et al*., 2013, Withey *et al*., 2020, Miller *et al*., 2017). Cocaine self-administration was then conducted for 5–9 weeks using the same 3-h session schedule.

### 2.4 Plasma collection and cocaine metabolite analysis

To validate oral cocaine intake, blood was collected approximately 24 h after the final cocaine self-administration session. Plasma was separated by centrifugation and stored at −80 °C until analysis. Cocaine and benzoylecgonine were measured using liquid chromatography mass spectrometry (LC-MS/MS) with a Sciex API 4000 system coupled to an Agilent 1100 HPLC system. Detailed chromatographic and MS/MS conditions are provided in Method S1 (Wansaw *et al*., 2005, Skopp *et al*., 2001, Kratz *et al*., 2021, Cognard *et al*., 2005).

### 2.5 ^18^F-FP-CIT PET imaging

To assess in-vivo DAT-related PET signals, 18F-FP-CIT PET imaging was performed before cocaine self-administration and 24 h after the final cocaine self-administration session. After intravenous injection of ^18^F-FP-CIT (123 kBq/g) through the tail vein, animals were allowed a 60-min uptake period, followed by 30-min PET acquisition using a Nanoscan positron emission tomography-computed tomography (PET-CT) system. DAT-related image signals were extracted from transverse PET images and normalized to body weight. Detailed PET reconstruction and semi-quantitative image analysis procedures are provided in Method S2.

### 2.6 MN9D cell culture and DAT internalization assay

MN9D dopaminergic cells (#SCC281; Sigma-Aldrich, St. Louis, MO, USA) were seeded at 3 × 10^6^ cells per 100-mm dish and differentiated for 9 days in expansion medium containing 1 mM N-butyrate. Differentiated cells were treated with 10-μM cocaine for 0, 10, 20, 30, or 60 min to examine DAT trafficking. Cells were harvested, and membrane and cytosolic fractions were separated using a commercial membrane protein extraction kit.

Protein concentrations were determined using the Bradford assay. Equal amounts of protein (50 μg) were separated by 12% sodium dodecyl sulfate–polyacrylamide gel electrophoresis (SDS-PAGE), transferred to polyvinylidene fluoride (PVDF) membranes, and incubated with antibodies against DAT (1:500; ab184451, Abcam, Cambridge, UK), E-cadherin (1:1,000; #3195, Cell Signaling Technology, Danvers, MA, USA), and GAPDH (1:2,000; 2118S, Cell Signaling Technology, Danvers, MA, USA). Membranes were incubated with HRP-conjugated secondary antibodies and visualized using chemiluminescence. Full uncropped blot images are provided in the Supplementary Information Fig. S2.

### 2.7 Ex-vivo brain slice imaging of DAT and GPR55 signals

Brains from drug-naïve marmosets were coronally sliced at 0.3-mm intervals to include the caudate nucleus, putamen, and nucleus accumbens. Brain slices were washed twice with Hank’s balanced salt solution for 30 min and treated with saline or cocaine (10 μM) for 30 min. After washing three times with phosphate buffered saline (PBS), slices were incubated with either fluorescent false neurotransmitter FFN-102 (10 μM; Sigma-Aldrich, St. Louis, MO, USA) to assess functional DAT-associated signal (Rodriguez *et al*., 2013) or T1117 (200 nM; Bio-Techne Corporation, Minneapolis, MN, USA) to assess GPR55-associated signal (Daly *et al*., 2010) for 1 h in the dark with gentle shaking. Fluorescence was measured using a VISQUE® in Vivo Smart-LF optical imaging system. Regions of interest corresponding to the caudate nucleus, putamen, and nucleus accumbens were defined according to the marmoset brain atlas; fluorescence intensity was quantified using CleVue software.

### 2.8 2-AG pretreatment during cocaine self-administration

To examine the effect of 2-AG on cocaine-taking behavior, marmosets were provided cocaine self-administration after baseline stable responding was established. Stability was defined as three consecutive sessions in which the number of cocaine deliveries did not vary by more than 25% of the mean number of rewards (Kohut and Bergman, 2016, Mello *et al*., 2013, Withey *et al*., 2020, Miller *et al*., 2017).

Marmosets received vehicle (2.5% DMSO and 2.5% Tween 80 in saline) or 2-AG (0.01 or 0.1 mg/kg, i.p.) 1 h before the cocaine self-administration session. The 2-AG doses were selected based on preliminary mouse locomotor activity data and interspecies dose conversion. Cocaine was delivered orally at 0.4 mg/mL, with 0.1 mL per active lever response. Active lever responses were analyzed across sessions, and cumulative response curves were plotted as the percentage of cumulative active lever responses over time from the start of the session.

### 2.9 Plasma 2-AG measurement

Plasma 2-AG levels were measured using ultra-high-performance liquid chromatography-high-resolution mass spectrometry (UHPLC-HRMS) with minor modifications of previously described methods (Pavón *et al*., 2013, Pedraz *et al*., 2015, Rivera *et al*., 2013a). Plasma samples collected in ethylenediamine tetraacetic acid (EDTA)-treated tubes were maintained at pH 5.8 on ice to reduce ex-vivo degradation of endocannabinoids. Lipids were extracted with tert-butyl methyl ether, dried under nitrogen, reconstituted, and analyzed using a Vanquish UHPLC system with high-resolution MS detection. Detailed UHPLC-HRMS conditions are provided in Supplementary Information Method S3.

### 2.10 Immunofluorescence analysis of GPR55 and DAT

Coronal marmoset brain sections (10-μm thick) were used for immunofluorescence analysis. Sections were incubated with primary antibodies against GPR55 (1:50; ab174700, Abcam, Cambridge, UK) and DAT (1:250; ab5990, Abcam, Cambridge, UK). After washing, sections were incubated with Alexa Fluor 488- and Alexa Fluor 568-conjugated secondary antibodies and DAPI (300 nM; Sigma-Aldrich, St. Louis, MO, USA). Images were acquired using an LSM 980 Axio Observer confocal microscope with Airyscan 2 (Carl Zeiss, Oberkochen, Germany) at × 200 magnification and analyzed using Zen Blue software (Carl Zeiss, Oberkochen, Germany).

### 2.11 Synaptosome preparation and dopamine quantification

Synaptosomes were isolated from marmoset brain regions, including the prefrontal cortex, striatum, and nucleus accumbens, using Syn-PER™ Synaptic Protein Extraction Reagent according to the manufacturer’s protocol. The resulting pellet was used as the synaptosomal fraction, and the supernatant was stored as the cytosolic fraction. Synaptosomal pellets were resuspended in artificial cerebrospinal fluid and incubated with artificial cerebrospinal fluid (ACSF) alone, 2-AG (8 ng/mL), cocaine (10 μM), or cocaine (10 μM) plus 2-AG (8 ng/mL) for 10 min at 37 °C. Dopamine levels in the supernatant were measured using HPLC with electrochemical detection. Detailed analytical methods are provided in Supplementary Information Method S4.

### 2.12 Western blot validation of synaptosome enrichment

Synaptosomal enrichment was validated via Western blotting using antibodies against synaptoporin and GAPDH. Synaptoporin was used as a synaptosomal marker, and GAPDH was used as a soluble/cytosolic marker. Detailed Western blot procedures are provided in Supplementary Information Method S5. Full uncropped blot images are provided in the Supplementary figure 2.

### 2.13 Statistical analysis

Data are presented as mean ± SEM. Active and inactive lever responses were analyzed using two-way analysis of variance (ANOVA) or two-way repeated-measures ANOVA followed by Holm–Sidak post-hoc tests. PET imaging values and fluorescence intensities from FFN-102 and T1117 imaging were analyzed using Student’s t-test. Cumulative active lever response curves were analyzed using the two-sample Kolmogorov–Smirnov test. Plasma 2-AG levels were analyzed using one-way ANOVA followed by Holm–Sidak post-hoc tests.

## 3. Results

### 3.1 Common marmosets acquired non-restraint oral cocaine self-administration

Oral cocaine self-administration was examined in four common marmosets under an FR1 schedule. The animals completed 24 self-administration sessions over 5–9 weeks under non-restraint, freely moving conditions. Individual animals in this cohort are referred to as SA1– SA4. Across the 24 sessions, group-level two-way repeated-measures ANOVA showed no significant main effect of session, lever, or session × lever interaction (session: F(23, 69) = 1.262, P = 0.227; lever: F(1, 3) = 5.164, P = 0.108; interaction: F(23, 69) = 1.548, P = 0.085; n = 4). However, Holm–Sidak multiple comparison analysis showed that the number of active lever presses were significantly greater than the number of inactive lever presses across the overall self-administration period (Fig. 1a; P = 0.023). Session-by-session comparisons showed that the number of active lever presses significantly higher than the number of inactive lever presses during sessions 7, 11, 20, 21, 22, and 24.

**Fig. 1.**
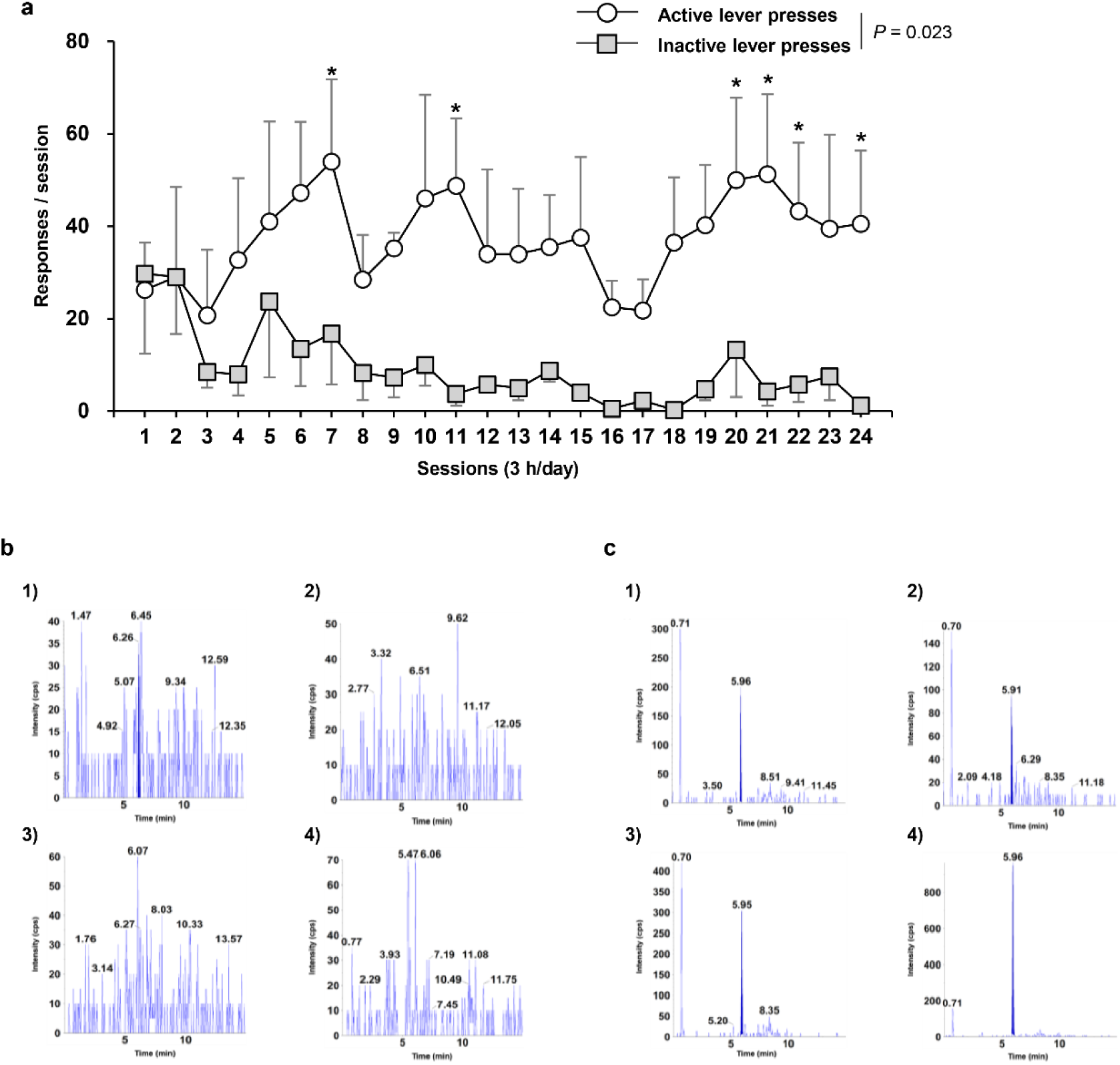
Lever-pressing behavior and LC-MS/MS analysis of plasma cocaine and benzoylecgonine in cocaine self-administering marmosets. (a) Active and inactive lever responses across 24 sessions under a fixed-ratio 1 (FR1) schedule of oral cocaine self-administration. Data are expressed as mean ± SEM (n = 4). Overall the number of active lever presses were significantly greater than the number of inactive lever presses across the entire self-administration period (P = 0.023). Session-by-session comparisons showed significantly higher active lever responses than inactive lever responses at selected session points. *P < 0.05 vs. inactive lever responses at the same session; two-way repeated-measures ANOVA followed by Holm–Sidak post-hoc test. (b) LC-MS/MS chromatograms for cocaine in marmoset plasma. The mass transition for cocaine was monitored from m/z 304.5 to 182.1, with a retention time of 6.04 min. Cocaine was not detected in plasma samples collected approximately 24 h after the final cocaine self-administration session from individual marmosets SA1–SA4. (c) LC-MS/MS chromatograms for benzoylecgonine, a major cocaine metabolite, in marmoset plasma. The mass transition for benzoylecgonine was monitored from m/z 290.1 to 105.1, with a retention time of 5.96 min. Distinct benzoylecgonine peaks were detected in plasma samples from all four marmosets SA1–SA4 after oral cocaine self-administration.

In addition, we summarized total cocaine intake, average lever responses, and intra-subject ANOVA results for each animal. All four marmosets showed a significant main effect of lever, with higher average active lever responses than inactive lever responses, although total cocaine intake varied among animals (Table 1). These results indicate preferential active lever responding during oral cocaine self-administration.

**Table 1.** Individual cocaine intake, lever responses, and intra-subject analysis during oral cocaine self-administration in common marmosets.

| Parameter | SA1 | SA2 | SA3 | SA4 |
| --- | --- | --- | --- | --- |
| Total cocaine intake (mg) | 56.24 | 47.54 | 67.30 | 142.50 |
| Average active lever responses | 25.08±3.44* | 10.83±2.31* | 40.33±2.96* | 73.08±4.56* |
| Average inactive lever responses | 8.21±2.82 | 5.17±1.44 | 13.29±3.18 | 10.29±3.95 |
| Session effect | F(23, 23) = 1.168, P = 0.356 | F(23, 23) = 1.746, P = 0.094 | F(23, 23) = 0.822, P = 0.679 | F(23, 23) = 1.229, P = 0.313 |
| Lever effect | F(1, 23) = 15.569, P < 0.001 | F(1, 23) = 5.961, P = 0.023 | F(1, 23) = 35.312, P < 0.001 | F(1, 23) = 120.765, P < 0.001 |

Data are expressed as mean ± SEM across all cocaine self-administration sessions for each animal. SA1–SA4 indicate individual animals in the cocaine self-administration cohort. Session and lever effects were analyzed by intra-subject two-way ANOVA. \*P < 0.05 vs. average inactive lever responses within the same animal.

To validate oral cocaine intake, plasma cocaine and benzoylecgonine were measured by LC-MS/MS approximately 24 h after the final cocaine self-administration session. Benzoylecgonine was detected in plasma, whereas cocaine itself was not detected (Table 2, Fig. 1b, c).

**Table 2.** Qualitative LC-MS/MS analysis of cocaine and benzoylecgonine in plasma after oral cocaine self-administration.

| Analyte | SA1 | SA2 | SA3 | SA4 |
| --- | --- | --- | --- | --- |
| Cocaine | - | - | - | - |
| Benzoylecgonine | + | + | + | + |

Plasma samples were collected approximately 24 h after the final oral cocaine self-administration session and subjected to qualitative LC-MS/MS analysis. SA1–SA4 indicate individual animals in the cocaine self-administration cohort. (+) detected; (–) not detected.

### 3.2 Cocaine self-administration reduced striatal DAT-related PET signal in vivo

To determine whether repeated oral cocaine self-administration altered DAT-related PET signal in vivo, ^18^F-FP-CIT PET imaging was performed before and after the cocaine self-administration period in the same animals as those used for behavioral acquisition analysis. Compared with the pre-self-administration baseline, marmosets showed a significant reduction in striatal DAT-related ^18^F-FP-CIT PET signals after repeated oral cocaine self-administration (Fig. 2a; P < 0.05, n = 4).

**Fig. 2.**
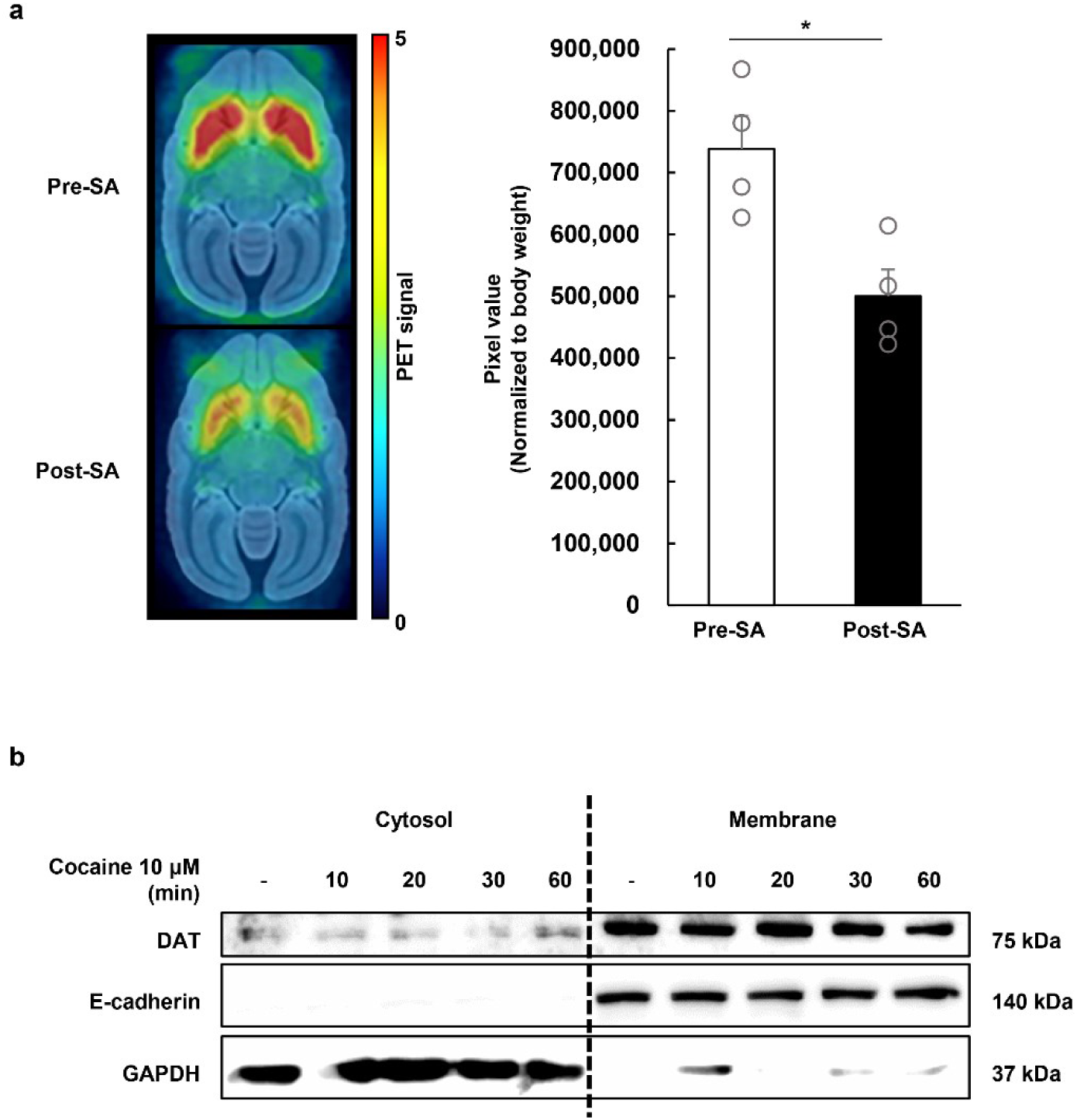
Cocaine self-administration reduced striatal dopamine transporter (DAT)-related positron emission tomography (PET) signals in vivo and altered DAT subcellular distribution in MN9D cells. (a) Representative ^18^F-N-(3-fluoropropyl)-2β-carboxymethoxy-3β-(4-iodophenyl) nortropane (18F-FP-CIT) PET images and semi-quantitative analysis of striatal DAT-related PET signals before and after oral cocaine self-administration in common marmosets. DAT-related image signals were quantified in the striatum using transverse PET images and normalized to body weight. Data are expressed as mean ± SEM (n = 4). *P < 0.05 vs. pre-self-administration; Student’s t-test. (b) Time-dependent effects of cocaine on DAT subcellular distribution in differentiated MN9D dopaminergic cells. Cells were treated with cocaine (10 μM) for 0, 10, 20, 30, or 60 min and subjected to subcellular fractionation into membrane and cytosolic fractions.

To examine whether cocaine exposure alters DAT subcellular distribution, differentiated MN9D dopaminergic cells were treated with cocaine and subjected to membrane/cytosolic fractionation. In this descriptive analysis, cocaine treatment produced an apparent time-dependent change in DAT distribution, with changes in cytosolic and membrane-associated DAT signals over time (Fig. 2b). Although this cell-based observation requires further quantitative replication, it provides a trend for cocaine-associated changes in DAT-related signals.

### 3.3 Cocaine altered DAT- and GPR55-associated fluorescent signals in marmoset striatal slices

To assess local neurochemical changes in marmoset striatal tissue, ex-vivo fluorescent ligand imaging was performed in coronal brain slices containing the caudate nucleus, putamen, and nucleus accumbens core. Functional DAT-associated signals were assessed using the fluorescent false neurotransmitter FFN-102. Quantification was performed using atlas-guided regions of interests (ROIs) placed within the caudate nucleus, putamen, and nucleus accumbens core. Acute cocaine treatment (10 μM, 30 min) reduced FFN-102 fluorescence intensity compared with saline control treatment (Fig. 3a; P < 0.05, n = 4), indicating decreased functional DAT-associated signals in marmoset striatal tissue.

**Fig. 3.**
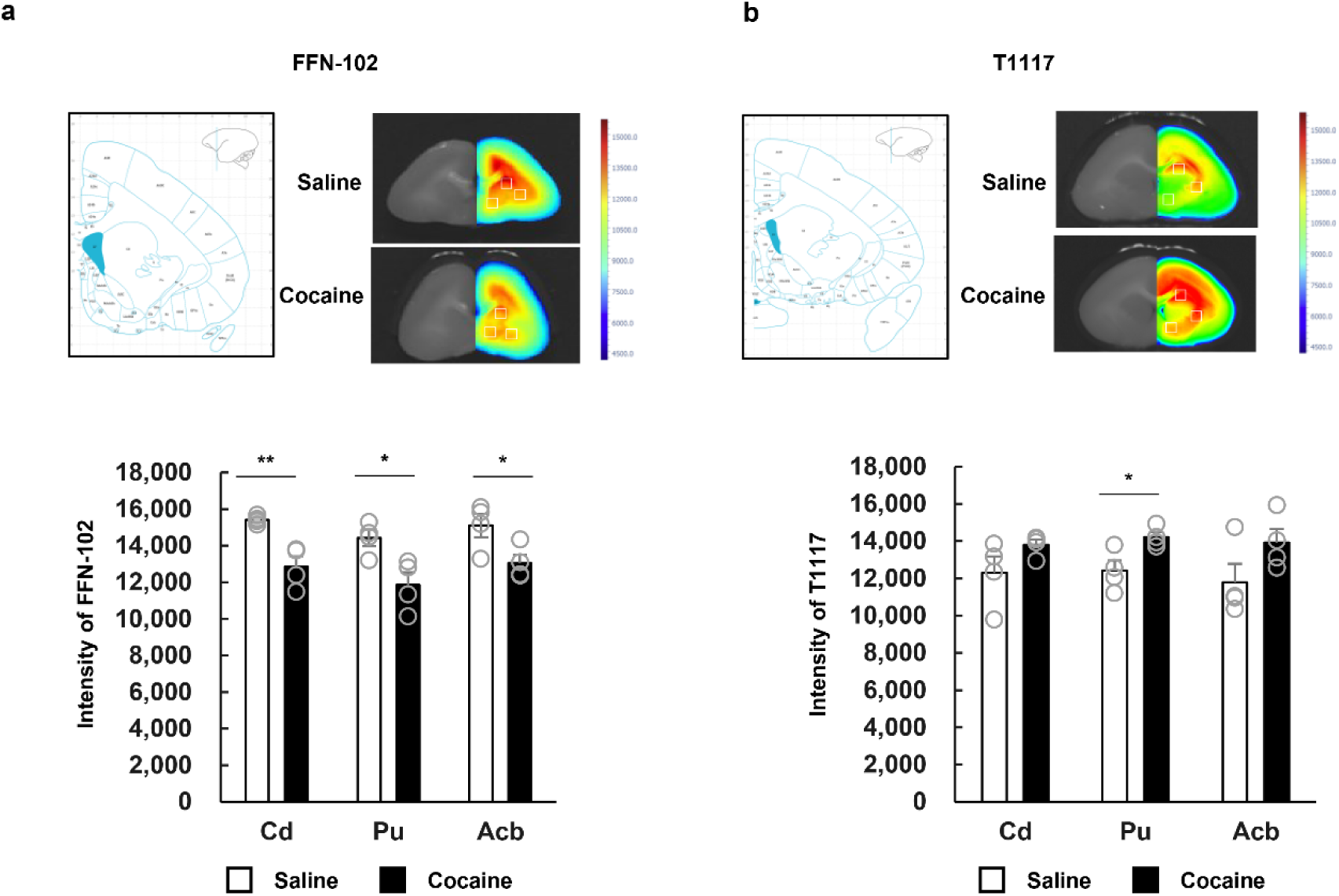
Acute cocaine treatment decreased fluorescent false neurotransmitter 102 (FFN-102)-associated DAT signals and increased T1117-associated GPR55 signals in marmoset striatal slices. (a) Representative coronal marmoset brain slice images and fluorescence heatmaps showing FFN102-associated DAT signals after saline or cocaine treatment. Brain slices containing the caudate nucleus (Cd), putamen (Pu), and nucleus accumbens (Acb) were treated with saline or cocaine (10 μM, 30 min), followed by incubation with FFN102. Fluorescence intensity was quantified using the same-sized rectangular ROIs placed within atlas-defined Cd, Pu, and Acb regions. (b) Representative coronal marmoset brain slice images and fluorescence heatmaps showing T1117-associated GPR55 signals after saline or cocaine treatment. Brain slices were treated with saline or cocaine (10 μM, 30 min), followed by incubation with T1117. Fluorescence intensity was quantified using the same-sized rectangular ROIs placed within atlas-defined Cd, Pu, and Acb regions. Warmer colors in the heatmaps indicate higher fluorescence intensity. Data are expressed as mean ± SEM (n = 4). *P < 0.05, **P < 0.01 vs. saline-treated control; unpaired Student’s t-test.

GPR55-associated signals were assessed using the fluorescent ligand T1117 (Daly *et al*., 2010). Cocaine treatment increased T1117 fluorescence intensity compared with saline control treatment. The increase was most evident in the putamen, whereas the caudate nucleus and nucleus accumbens core showed a similar increasing trend (Fig. 3b; P < 0.05, n = 4). These results suggest that cocaine exposure decreases DAT-associated functional signals while increasing T1117-associated GPR55 signals in marmoset striatal tissue.

### 3.4 2-AG pretreatment suppressed cocaine-taking behavior and restored circulating 2-AG levels

To determine whether 2-AG changes cocaine-taking behavior, systemic 2-AG pretreatment was examined in a separate cohort of marmosets under the oral cocaine self-administration paradigm. Individual animals in this cohort are referred to as AG1–AG3. Marmosets received vehicle or 2-AG (0.01 or 0.1 mg/kg, i.p.) 1 h before cocaine self-administration.

Compared with the basal cocaine self-administration condition, 2-AG pretreatment reduced active lever responding for cocaine (Fig. 4a). Two-way repeated-measures ANOVA showed a significant main effect of lever but not session or session × lever interaction (session: F(2, 20) = 1.396, P = 0.252; lever: F(1, 2) = 40.867, P = 0.024; interaction: F(10, 20) = 1.510, P = 0.207; n = 3). Cumulative event-time analysis showed a leftward shift in the distribution of active lever responses toward the earlier part of the session after 2-AG pretreatment compared with vehicle pretreatment (Fig. 4b; P < 0.05, two-sample Kolmogorov–Smirnov test). This pattern shows that the remaining active responses occur earlier in the session, with less responding during the later phase of the session.

**Fig. 4.**
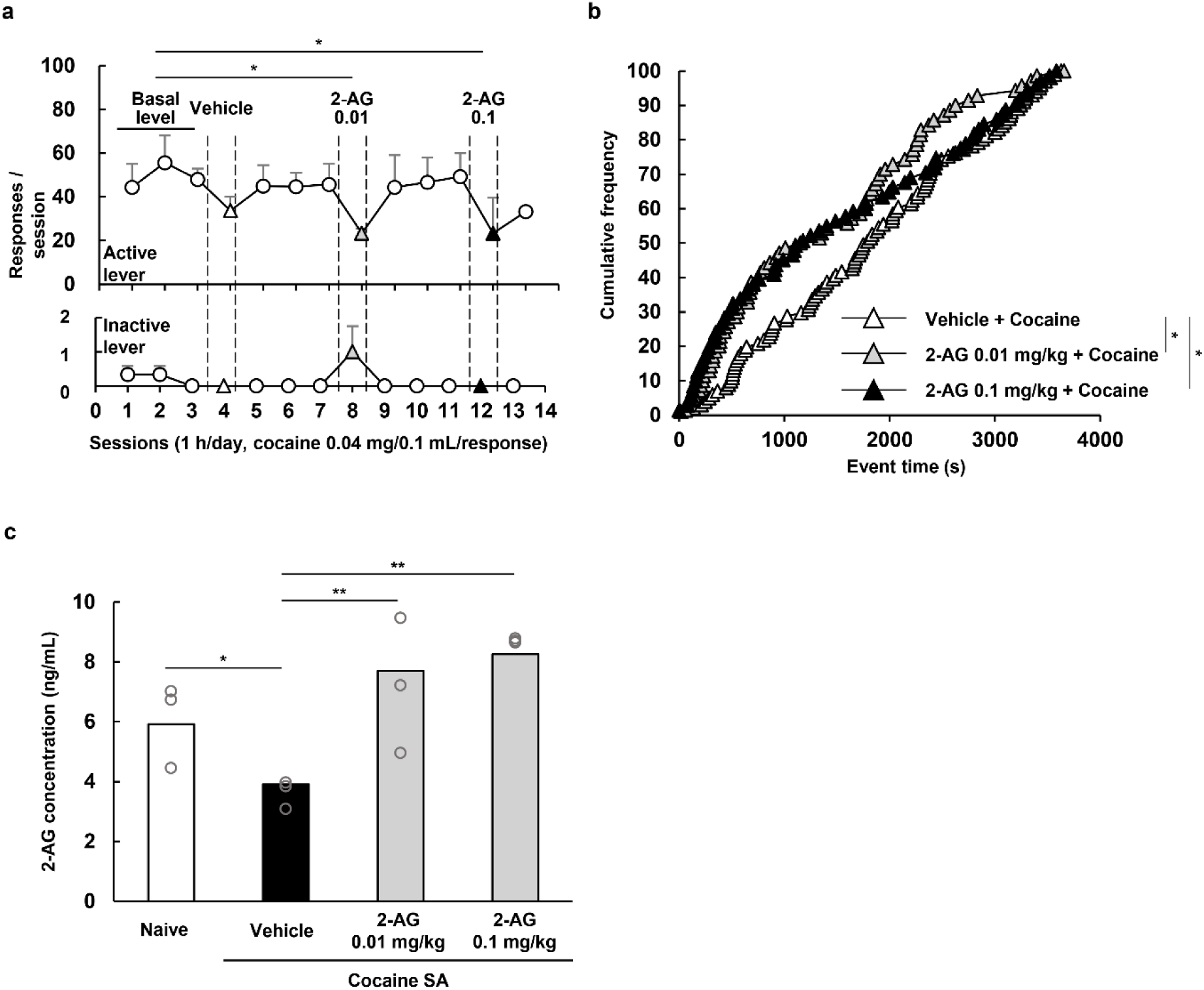
Effects of 2-AG pretreatment on cocaine self-administration and plasma 2-AG levels in common marmosets. (a) Effects of 2-arachidonoylglycerol (2-AG) pretreatment on active and inactive lever responses under a fixed-ratio 1 (FR1) schedule of oral cocaine self-administration. Marmosets were administered vehicle (2.5% DMSO and 2.5% Tween 80 in saline, 2 mL/kg, i.p.) or 2-AG (0.01 or 0.1 mg/kg, i.p.) 1 h before cocaine self-administration testing. Cocaine was delivered orally at 0.4 mg/mL and 0.1 mL/response. Data are expressed as mean ± SEM (n = 3). *P < 0.05 vs. basal level; two-way repeated-measures ANOVA followed by Holm–Sidak post-hoc test. (b) Cumulative frequency distribution of active lever response event times during cocaine self-administration after vehicle or 2-AG pretreatment. A leftward shift of the cumulative event-time distribution toward earlier time points indicates that active responses occurred earlier within the session. Differences in event-time distributions were analyzed using the two-sample Kolmogorov–Smirnov test. *P < 0.05 vs. vehicle. (c) Plasma concentrations of 2-AG in drug-naïve and cocaine self-administering marmosets after pretreatment with vehicle or 2-AG (0.01 or 0.1 mg/kg). Data are expressed as mean ± SEM (n = 3). *P < 0.05, **P < 0.01; one-way ANOVA followed by Holm–Sidak post-hoc test.

To assess whether cocaine self-administration was associated with changes in circulating endocannabinoid levels, plasma 2-AG concentrations were measured by UHPLC-HRMS. Plasma 2-AG levels differed significantly among groups (Fig. 4c; F(3, 8) = 7.55, P < 0.001; n = 3). Post-hoc analysis showed that plasma 2-AG concentrations were significantly lower in the cocaine self-administration vehicle-treated group than in the drug-naïve control group (Fig. 4c; P < 0.05). Pretreatment with 2-AG at both 0.01 and 0.1 mg/kg restored plasma 2-AG levels after cocaine self-administration (Figure 4c; P < 0.01). These results show that systemic 2-AG pretreatment reduces cocaine-taking behavior and restores the cocaine-associated decrease in circulating 2-AG levels.

### 3.5 GPR55 and DAT showed spatially associated signals, and 2-AG enhanced cocaine-induced dopamine elevation in marmoset synaptosomes

To examine the anatomical relationship between GPR55 and DAT in marmoset brain tissue, confocal immunofluorescence imaging was performed using antibodies against GPR55 and DAT. GPR55-positive and DAT-positive signals were detected in the same brain sections. Merged images showed colocalized GPR55 and DAT immunofluorescent signals in marmoset brain sections (Fig. 5a). These findings suggest that GPR55-positive and DAT-positive signals are present in the same anatomical regions.

**Fig. 5.**
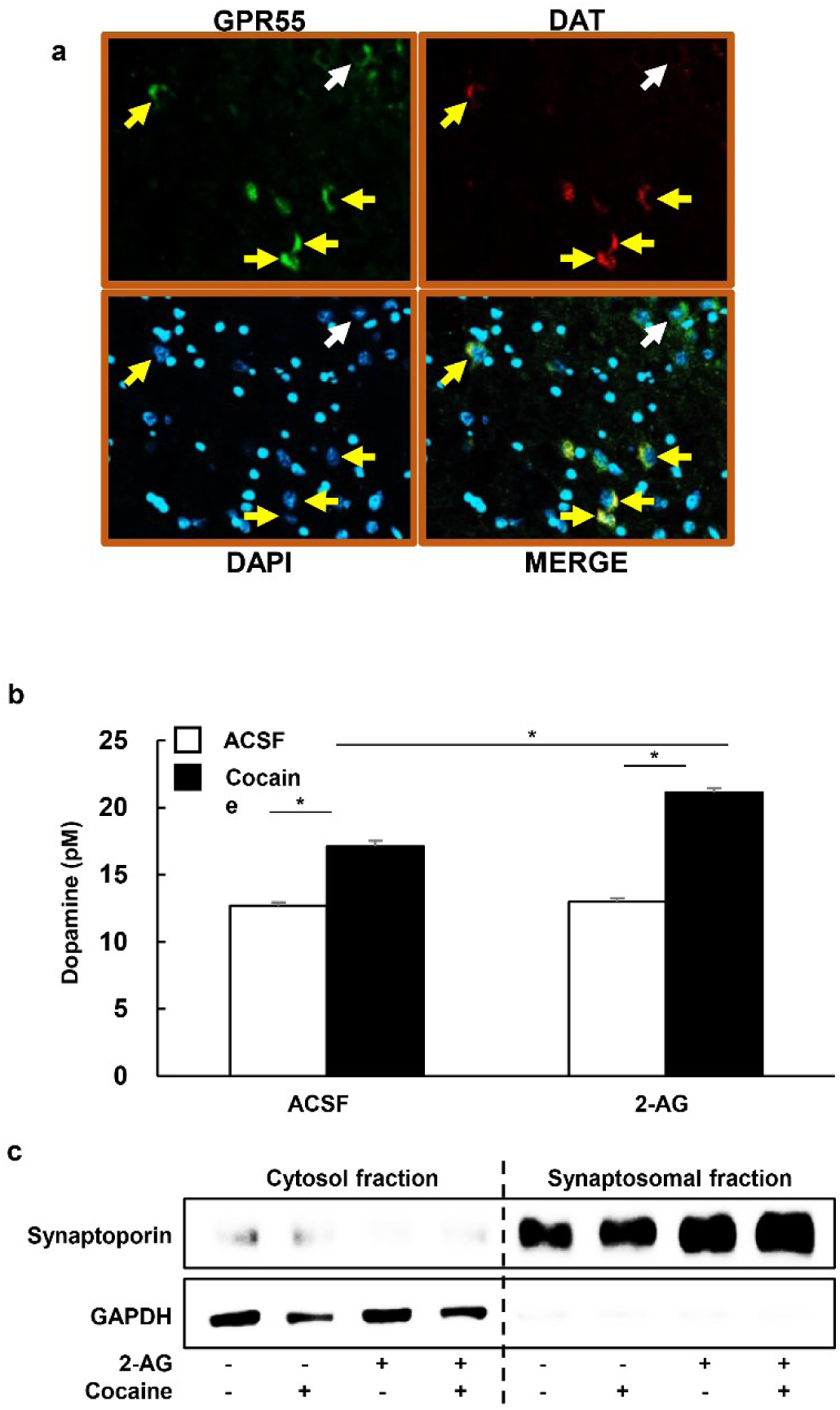
Spatial association of G protein-coupled receptor 55 (GPR55) and DAT signals and 2-AG-enhanced dopamine elevation in marmoset synaptosomes. (a) Representative confocal immunofluorescence images showing GPR55-positive and DAT-positive signals in marmoset brain sections. Sections were immunostained for GPR55 (green) and DAT (red) and counterstained with DAPI (blue). Yellow arrows indicate cells or regions showing partially merged GPR55/DAT signals in the merged image, whereas the white arrow indicates a representative region without apparent merged GPR55/DAT signal. (b) Dopamine levels in the supernatant of purified marmoset synaptosomal preparations. Synaptosomes were incubated with artificial cerebrospinal fluid (ACSF), 2-AG (8 ng/mL), cocaine (10 μM), or cocaine plus 2-AG. Data are expressed as mean ± SEM (n = 3). *P < 0.05; two-way ANOVA followed by Holm–Sidak post-hoc test. (c) Western blot validation of synaptosomal fraction enrichment. Synaptosomal enrichment was confirmed by increased synaptoporin signals in the synaptosomal pellet fraction. GAPDH, used as a soluble/cytosolic marker, was detected mainly in the cytosolic fraction and was reduced in the synaptosomal fraction. Treatment with 2-AG and/or cocaine did not markedly alter the fractionation profile.

To assess dopamine responses at the isolated synaptic level, synaptosomes were prepared from marmoset brain tissue. Western blotting confirmed enrichment of the synaptosomal fraction. Synaptoporin, a synaptic vesicle-associated protein, was enriched in the synaptosomal pellet fraction, whereas GAPDH, used as a soluble/cytosolic marker, was detected mainly in the cytosolic fraction and reduced in the synaptosomal fraction (Fig. 5c).

Dopamine levels in the synaptosomal supernatant were measured after incubation with ACSF, 2-AG, cocaine, or cocaine plus 2-AG. Cocaine (10 μM) increased dopamine levels in the synaptosomal preparation, consistent with blockade of dopamine reuptake (Fig. 5b). Co-incubation with 2-AG (8 ng/mL) further enhanced cocaine-induced dopamine elevation compared with cocaine treatment alone (Fig. 5b; P < 0.05, n = 3). These results suggest that 2-AG can increase cocaine-induced dopaminergic responses in marmoset synaptosomal preparations.

## 4. Discussion

In this study, we established a non-restraint oral cocaine self-administration model in freely moving common marmosets and tested whether systemic 2-AG pretreatment reduces cocaine-taking behavior. In addition, we examined cocaine-associated changes in DAT-related PET signals, GPR55-associated signals, circulating 2-AG levels, and synaptosomal dopamine responses.

Common marmosets acquired oral cocaine self-administration under an FR1 schedule, as shown by preferential active lever responding. Cocaine self-administration has been widely used in non-human primates to model drug reinforcement and cocaine-taking behavior (Kohut *et al*., 2020, Mandeville *et al*., 2011, Porter *et al*., 2011). Many primate cocaine self-administration studies use intravenous catheter-based procedures, which allow precise drug delivery but require surgery and specialized housing. Cocaine can be used by several routes in humans, including oral, intravenous, intranasal, and inhalational routes (National Institute on Drug, 2016). Therefore, a non-restraint oral self-administration model in marmosets may be a useful additional model for studying cocaine reinforcement under freely moving conditions. In this study, oral cocaine intake was supported by active lever discrimination and by plasma detection of benzoylecgonine, the major stable metabolite of cocaine. Cocaine was not detected in plasma collected approximately 24 h after the final self-administration session, consistent with the relatively short half-life of cocaine in primates (Zhou *et al*., 2001, Parker and Laizure, 2010, Jufer *et al*., 1998). Thus, benzoylecgonine detection confirmed oral cocaine ingestion during the self-administration procedure.

Repeated oral cocaine self-administration was associated with reduced striatal DAT-related ^18^F-FP-CIT PET signal in vivo. Cocaine binds to DAT and can alter transporter function and conformation (Xu and Chen, 2020). PET imaging was performed after the self-administration period, when parent cocaine was not detected in plasma. Therefore, the reduced PET signals may reflect longer-lasting cocaine-associated changes in DAT-related tracer binding rather than acute competition by circulating cocaine. This interpretation is consistent with reports of cocaine-related changes in dopamine transporter function and regulation (Harraz *et al*., 2022, Gabrielsen *et al*., 2011, Cagniard *et al*., 2014). Ex-vivo fluorescent ligand imaging supported the PET results, as acute cocaine treatment decreased FFN-102 signals in marmoset striatal slices, indicating reduced functional DAT-associated signals. In differentiated MN9D dopaminergic cells, cocaine exposure produced a descriptive time-dependent change in membrane-associated and cytosolic DAT signals. This result suggests a preliminary trend toward cocaine-associated changes in DAT-related signals.

Furthermore, the present study found that cocaine exposure increased T1117-associated signals in marmoset striatal slices, most clearly in the putamen, while reducing FFN-102-associated DAT signals. Cocaine can regulate 2-AG synthesis and release in the midbrain, and endocannabinoid signaling contributes to cocaine-related neuroadaptations (Wang *et al*., 2015, Nakamura *et al*., 2019, Engi *et al*., 2021, Zhang *et al*., 2015, Adamczyk *et al*., 2012). Moreover, CB1 and CB2 receptor-related mechanisms have been implicated in cocaine reward and cocaine-related behaviors (Vlachou *et al*., 2003, Fattore *et al*., 1999, Xi *et al*., 2011). In this study, animals with cocaine self-administration showed reduced circulating 2-AG levels compared with drug-naïve controls, consistent with reports of altered circulating endocannabinoid levels in substance use conditions (González *et al*., 2002, Bystrowska *et al*., 2014). These results suggest that cocaine self-administration is associated with both dopaminergic and endocannabinoid-related changes in marmosets.

A key finding of this study is that systemic 2-AG pretreatment reduces cocaine-taking behavior. No approved pharmacotherapy is currently available for cocaine use disorder, and clinical studies of cannabinoid-related approaches, including cannabidiol, have not consistently shown efficacy against cocaine craving or relapse (Daldegan-Bueno *et al*., 2021, Mongeau-Pérusse *et al*., 2021, Meneses-Gaya *et al*., 2021). Previous human studies reported reduced 2-AG levels in abstinent participants (Lu *et al*., 2019, Pavón *et al*., 2013, Pedraz *et al*., 2015), and experimental studies have suggested that 2-AG signaling can affect cocaine-induced motivation and dopaminergic plasticity (Wang *et al*., 2015, Nakamura *et al*., 2019, Engi *et al*., 2021, Wang *et al*., 2020). In the present marmoset model, 2-AG pretreatment at 0.01 and 0.1 mg/kg reduced active lever responding for cocaine and restored the cocaine-associated decrease in circulating 2-AG levels. These results support the idea that 2-AG signaling acts as an endogenous regulator of cocaine reinforcement.

The leftward shift in cumulative response timing toward the earlier part of the session, together with the reduction in total active lever responding, suggests that 2-AG promotes earlier termination of cocaine-taking behavior after initial responding. This satiety-like pattern may be related to the synaptosome finding that 2-AG enhanced cocaine-induced dopamine elevation in marmoset synaptosomal preparations. The 2-AG concentration used in this ex-vivo assay was selected to reflect plasma concentrations measured in vivo. These findings suggest that 2-AG strengthens the early dopaminergic response to cocaine and reduces later responding during the self-administration session.

Additionally, confocal immunofluorescence showed spatial association between GPR55-positive and DAT-positive signals in marmoset brain sections. Together with the cocaine-induced increase in T1117-associated signal, these results suggest that 2-AG influences cocaine-related dopamine dynamics through a GPR55/DAT-associated dopaminergic mechanism. This interpretation is consistent with the behavioral finding that 2-AG reduced total active lever responding and shifted the remaining responses toward earlier time points within the session. Thus, 2-AG may regulate cocaine reinforcement by changing the timing and strength of dopaminergic responses after initial cocaine exposure.

The complex roles of GPR55 in drug reward have been reported in previous studies. For example, GPR55 activation inhibited nicotine- or amphetamine-induced conditioned place preference, whereas the GPR55 agonist O-1602 did not block cocaine self-administration in mice (Liu *et al*., 2021, Sánchez-Zavaleta *et al*., 2023, Xi *et al*., 2024). Together with our findings, this suggest that the behavioral effects of 2-AG in this marmoset model may involve multiple endocannabinoid-related targets and downstream pathways.

In addition, another endogenous cannabinoid such as 2-LG has partial agonist properties at CB1, and its pharmacological profile may differ from that of 2-AG (Lu *et al*., 2019, Mechoulam *et al*., 1995, Veilleux *et al*., 2019). Therefore, elucidation of the effects of various endocannabinoids remains an important objective for future research. Other enzymatic and neuronal pathways should also be considered. Thus, 2-AG levels are regulated by metabolic enzymes, including monoacylglycerol lipase (MAGL), ABHD6, and ABHD12 (Veilleux *et al*., 2019, Trigo and Le Foll, 2016, Senior and Isselbacher, 1963, Vaughan *et al*., 1964, Deng and Li, 2020). Because MAGL and related enzymes can regulate 2-AG and other lipid metabolites, future studies should examine whether changes in 2-AG metabolism contribute to cocaine-taking behavior in marmosets. Non-cannabinoid pathways may also participate in the behavioral effects of 2-AG. For example, 5-HT1A receptor signaling is associated with cocaine-seeking behavior, and cannabinoid-related compounds can interact with serotonergic systems (Oakes *et al*., 2019, You *et al*., 2016, Katsidoni *et al*., 2013). Therefore, the effect of 2-AG on cocaine self-administration may reflect interactions among endocannabinoid, dopaminergic, and other neuromodulatory systems.

This study has several limitations. First, the number of marmosets was small (n = 4). Second, the role of GPR55 was inferred from associated signals and synaptosomal dopamine responses, without receptor blockade or genetic manipulation. Third, direct evidence for reward satiety will require within-session measurements of response rate, inter-response interval, and dopamine in vivo. Finally, the MN9D DAT trafficking experiment was descriptive and did not allow the direct examination of 2-AG effects.

In conclusion, this study supports common marmosets as a useful primate model for studying cocaine reinforcement and endocannabinoid regulation. Oral cocaine self-administration was associated with reduced striatal DAT-related PET signals and decreased circulating 2-AG levels. Systemic 2-AG pretreatment reduced cocaine-taking behavior and restored circulating 2-AG levels, while a possible dopamine-related mechanism was suggested by DAT/GPR55-associated assays. These findings support further study of endocannabinoid-dopamine interactions in primate models of cocaine reinforcement.

## Acknowledgments

This work was supported by the Ministry of Food and Drug Safety of the Republic of Korea (20182MFDS425, 22214MFDS252, and 23212MFDS217); the Bio and Medical Technology Development Program of the National Research Foundation of Korea (NRF), funded by the Korean government (MSIT) (RS-2024-00440787); the NRF grant funded by the Korean government (MSIT) through the Medical Research Center Program (RS-2025-02273102), funded by the Ministry of Education (2021RIS-001); the Basic Science Research Program through the NRF, funded by the Ministry of Education (RS-2024-00460411 and NRF-2021R1I1A1A01058188); a 2023 grant from the Korean Society of Ginseng; and the Chungbuk National University BK21 program (2023), and the Regional Innovation System & Education(RISE) program through the (Chungbuk Regional Innovation System & Education Center), funded by the Ministry of Education(MOE) and the (Chungcheongbuk-do), Republic of Korea. (2026-RISE-11-014-03)

## Author contributions

CKL, HE, TY, DL, and JY designed the study. SMG, HE, TY, JJL, HK, SMG, SBN, EK, THK, CWP, BYH, and SSY performed the experiments and contributed to data acquisition. CKL, HE, TY, HK, DL, and JY analyzed and interpreted the data. CKL, HE, TY, DL, and JY wrote and revised the manuscript. DL and JY supervised the study. All authors read and approved the final manuscript.

## Declaration of competing interest

The authors declare no competing interests.

## Supplementary materials

Supplementary material associated with this article can be found, in the online version

## Supporting Information

### 1. Supporting methods

#### Method S1: LC-MS/MS Analysis for Cocaine and Benzoylecgonine

Cocaine and benzoylecgonine in marmoset plasma were analyzed using an API 4000 LC-MS/MS system coupled to an Agilent 1100 HPLC system. Data acquisition and peak integration were performed using Analyst 1.6.3 software. Chromatographic separation was performed on a Kinetex C18 100 Å column (2.1 × 50 mm, 2.6 μm).

The mobile phase consisted of 0.1% formic acid in distilled water (A) and methanol (B). The flow rate was 0.2 mL/min, the column oven temperature was maintained at 40°C, and the injection volume was 10 μL. The gradient program was as follows.

| Time (min) | A (%) | B (%) |
| --- | --- | --- |
| 0.0 | 90 | 10 |
| 1.5 | 90 | 10 |
| 3.0 | 5 | 95 |
| 5.0 | 5 | 95 |
| 8.0 | 90 | 10 |
| 15.0 | 90 | 10 |

MS/MS analysis was performed using electrospray ionization in multiple reaction monitoring mode with positive polarity. The MRM transitions were selected based on previously reported methods and optimized for cocaine and benzoylecgonine.

| Analyte | Q1 | Q3 | DP (V) | CE (V) | CXP (V) |
| --- | --- | --- | --- | --- | --- |
| Cocaine | 304.5 | 182.1 | 56 | 27 | 12 |
| Benzoylecgonine | 290.1 | 105.1 | 60 | 28 | 13 |

#### Methods S2: PET image reconstruction and semi-quantitative analysis

PET images were reconstructed using a Monte Carlo-based Tera-Tomo 3D reconstruction algorithm with attenuation and scatter correction. Striatal and cerebellar regions of interest were defined using PMOD 3.8. For the semi-quantitative analysis shown in Fig. 2A, transverse PET images were exported and analyzed using RadiAnt DICOM Viewer and ImageJ. DAT-related image signals were extracted from the striatal region of interest and normalized to body weight. The resulting values were used as semi-quantitative indices of striatal DAT-related ¹⁸F-FP-CIT PET signal.

#### Method S3: UHPLC-HRMS Analysis for 2-AG

Plasma 2-arachidonoylglycerol (2-AG) levels were measured using UHPLC-HRMS. Plasma samples collected in EDTA-treated tubes were kept on ice, and the pH was maintained at 5.8 during processing to reduce ex vivo degradation or acyl migration of endocannabinoids. Plasma was diluted 1:1 with distilled water and extracted with tert-butyl methyl ether at a 1:6 ratio. After centrifugation at 2,800 × g for 5 min at room temperature, the organic phase from 100 μL plasma was collected, evaporated under nitrogen at 40°C, and reconstituted in water:acetonitrile (10:90) containing 0.1% formic acid.

Samples were analyzed using a Thermo Vanquish UHPLC system equipped with photodiode array and mass spectrometry detectors. Chromatographic separation was performed on a Unison UK-C18 MF column (2.0 × 50 mm, 3.0 μm). The mobile phase consisted of 0.1% formic acid in distilled water (A) and 0.1% formic acid in acetonitrile (B). The flow rate was 0.3 mL/min, the column oven temperature was maintained at 40°C, and the injection volume was 5 μL. The gradient program was as follows.

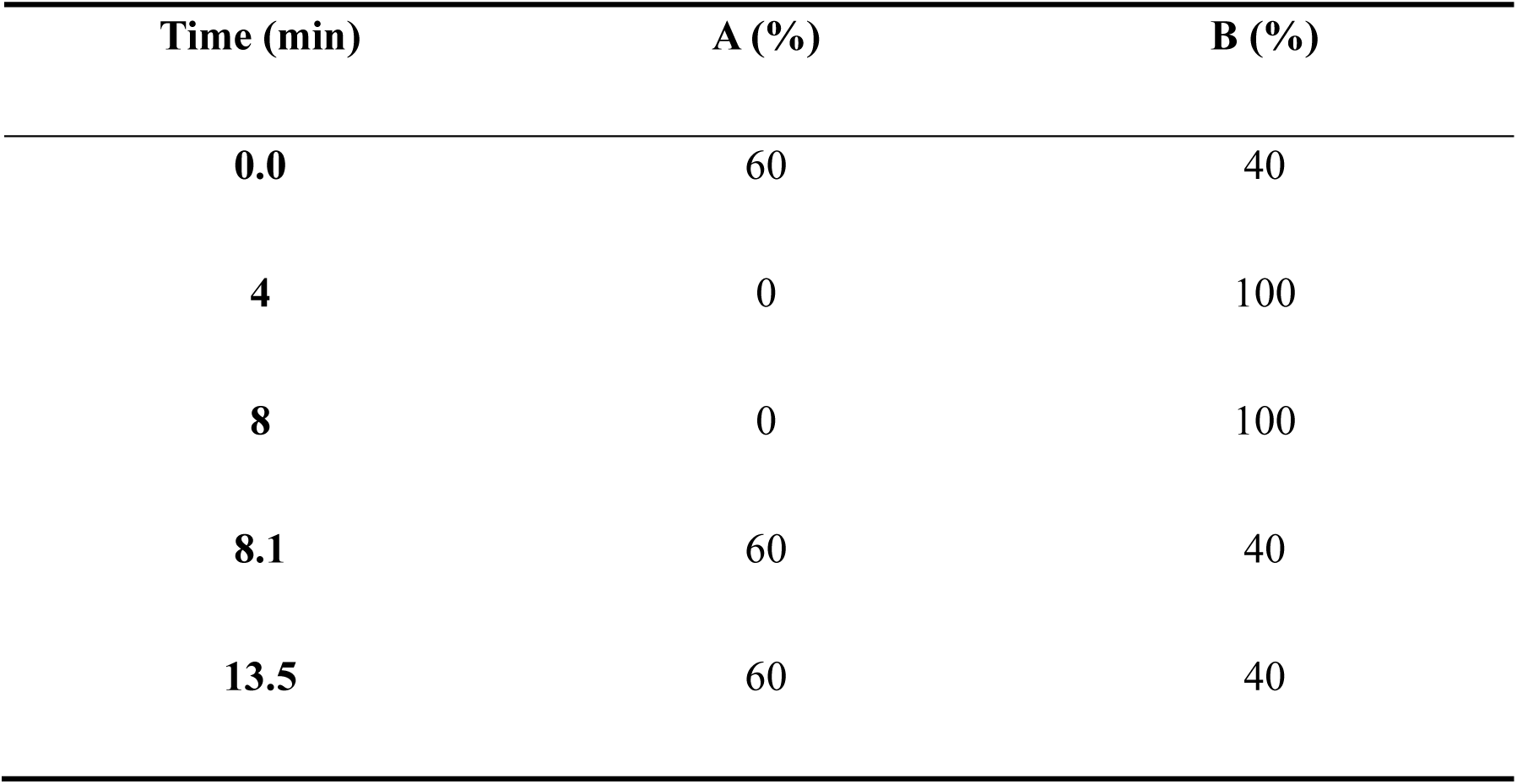

The HRMS parameters were as follows.

| Parameters | Conditions |
| --- | --- |
| Spray voltage (V) | Positive = 3,500, Negative = 2,500 |
| Gas (arb) | Sheath gas = 50, Aux gas = 15, Sweep gas = 1 |
| Temperature ($^{\circ}$C) | Ion transfer tube = 325, Vaporizer = 350 |
| Resolution | 60,000 |
| Scan range | 200-2,000 (m/z) |

Parallel reaction monitoring parameters for 2-AG were as follows.

| PRM (Parallel Reaction Monitoring) parameters |
| --- |

| Target | Molecular weight | Retention time (min) | Precursor ion (*m/z*) | Product ion (*m/z*) | Collision energy (eV) |
| --- | --- | --- | --- | --- | --- |
| 2-AG | 378.2770 | 4.54 | 379.2843 | 287.2379 | 15 |

#### Method S4. HPLC analysis of dopamine in synaptosomal supernatants

Dopamine levels in synaptosomal supernatants were measured using an HPLC system equipped with an electrochemical detector. Samples were injected onto an EiCOMPAK SC-5ODS column. The mobile phase consisted of 87% 0.1 M acetic acid-citric acid, 13% methanol, 50 mg/L sodium 1-octanesulfonate, and 5 mg/L EDTA·Na, pH 3.5, at a flow rate of 230 μL/min.

#### Methods S5: Western blot analysis for DAT fractionation and synaptosome validation

For the MN9D DAT fractionation experiment, membrane and cytosolic fractions were prepared using a commercial membrane protein extraction kit. Protein concentrations were determined by Bradford assay. Equal amounts of protein were separated by 12% SDS-PAGE and transferred to PVDF membranes. Membranes were incubated with antibodies against DAT, E-cadherin, and GAPDH. E-cadherin was used as a membrane-associated marker, and GAPDH was used as a cytosolic marker. HRP-conjugated secondary antibodies were applied, and immunoreactivity was visualized using chemiluminescence.

For synaptosome validation, synaptosomal pellets were lysed in RIPA buffer and incubated on ice for 60 min, followed by centrifugation at 13,000 rpm for 20 min at 4°C. Equal amounts of protein were separated by 12% SDS-PAGE and transferred to PVDF membranes. Membranes were incubated with antibodies against synaptoporin and GAPDH. Synaptoporin was used as a synaptosomal marker, and GAPDH was used as a soluble/cytosolic marker. Full uncropped blot images are provided in Supporting Information Fig. S3.

## Supplementary Figure Legends

**Fig. S1.**
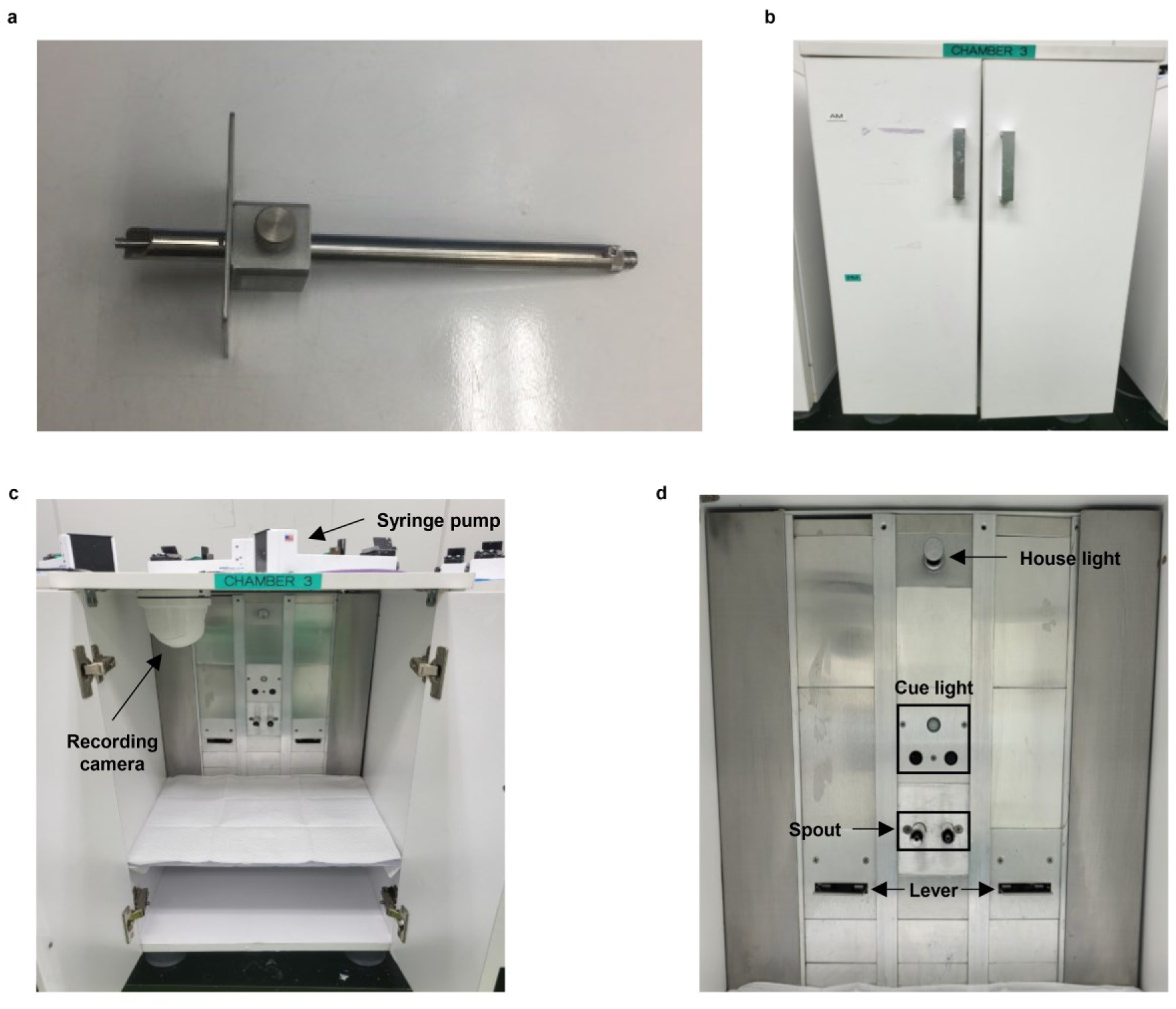
Design and configuration of the operant self-administration apparatus for common marmosets. (a) Close-up view of the fluid delivery spout. (b) External view of the custom operant chamber. (c) Internal components of the operant chamber. (d) Schematic layout of the self-administration assembly.

**Fig. S2.**
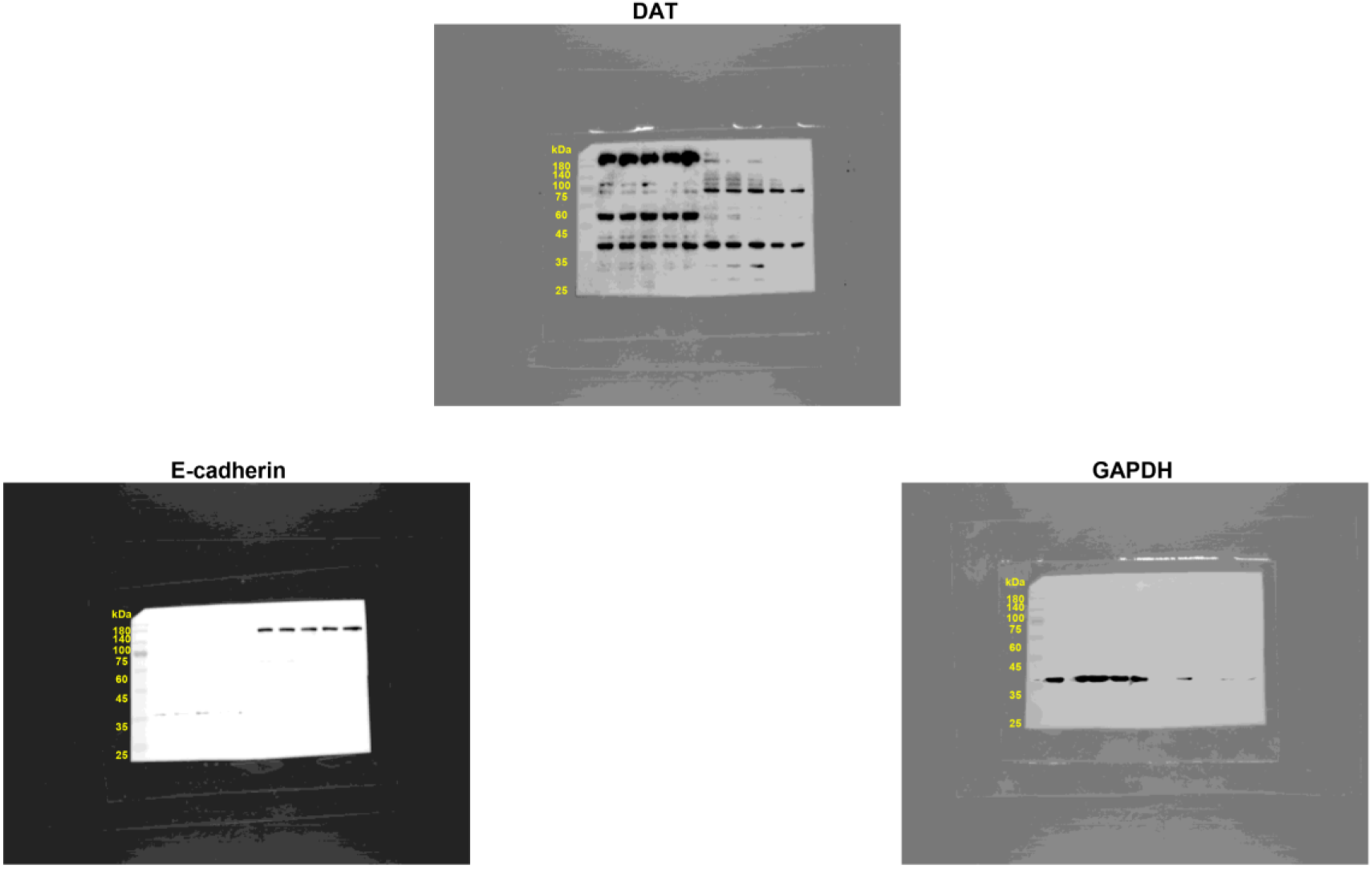
Full-length, uncropped western blot images for. Figure 2. Original immunoblot membranes showing fractionated protein levels of dopamine transporter (DAT), E-cadherin and GAPDH in the MN9D cells. Molecular weight markers (kDa) are indicated on the left of each blot.

## Supplementary Video Legends

**Video S1. Representative behavioral recording showing an active lever-press response during oral cocaine self-administration.** The unrestrained common marmoset interacted with the operant device, resulting in oral cocaine delivery after pressing the active lever.

**Video S2. Representative behavioral recording showing an inactive lever-press response during oral cocaine self-administration.** The common marmoset pressed the inactive lever, which recorded non-reinforced lever interaction.

## Notes

### Competing Interest Statement

The authors have declared no competing interest.

