## Supplementary figures and images for "Restoration of circulating 2-arachidonoylglycerol levels attenuates cocaine self-administration through endocannabinoid–dopamine interactions in common marmosets"

### Supplementary_Figure S1..tif

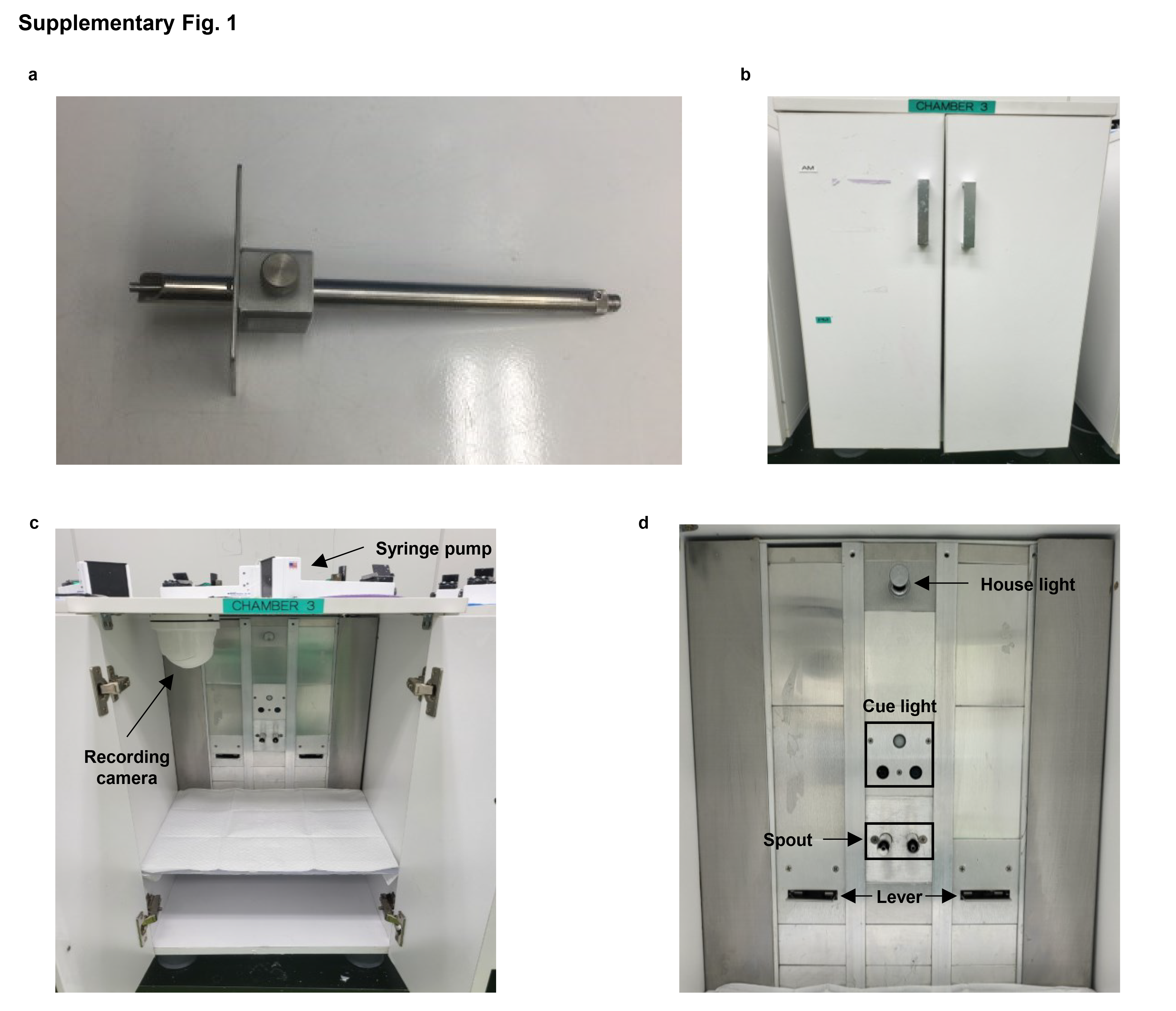

### Supplementary_Figure S2.tif

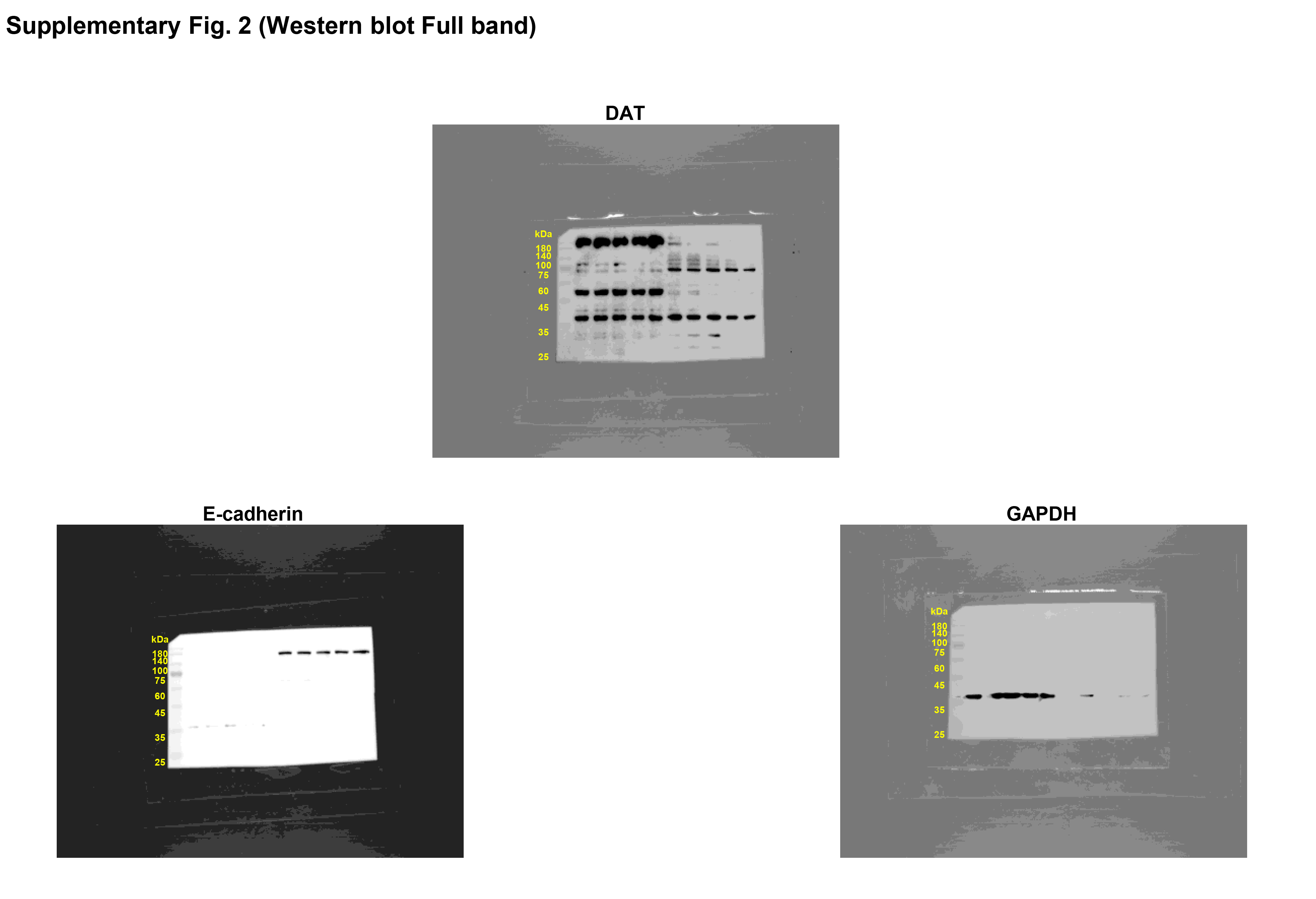
